# Computational mapping of antibody-receptor energy landscapes to predict internalization

**DOI:** 10.64898/2026.03.13.711720

**Authors:** Pablo Llombart, Cristina Nieto-Jiménez, Atanasio Pandiella, Alberto Ocana, Jorge R. Espinosa

## Abstract

Antibody internalization is critical for the action of antibody–drug conjugates, yet antibody discovery pipelines typically prioritize binding affinity rather than functional internalization. Here we show that molecular dynamics simulations can map the binding energy landscape between antibody clones and the membrane protein JAM-A, enabling computational predictions of antibody internalization. Using the sequences of newly generated anti–JAM-A monoclonal antibodies (mAbs), we perform atomistic potential-of-mean force simulations to evaluate the binding free energy to the JAM-A receptor and their interaction fingerprint at residue-level resolution. We find that internalizing mAb derived from different hybridomas exhibit a unique membrane-oriented contact topology that promotes cooperative receptor–receptor interactions, lowering the energetic barrier for early endocytic events. Reconstruction of the receptor binding energy landscape further reveals that electrostatic interactions between charged residues and multivalent cation-*π* and polar interactions correlate with successful mAb internalization in ovarian cancer cells. In contrast, strong binding affinity of the fragment antigen-binding domain correlates with poor internalization. Together, our results establish molecular dynamics–guided clonal selection as a predictive framework for optimizing internalizing therapeutic antibodies and provide mechanistic insight into how antibody binding reshapes membrane-proximal receptor energetics to drive endocytosis.

## 1. INTRODUCTION

Antibody–drug conjugates (ADCs) have reshaped the therapeutic landscape of oncology by enabling the intracellular delivery of cytotoxic agents to tumor cells^1,2^. Since the clinical introduction of Gemtuzumab ozogamicin and subsequent approvals such as Trastuzumab emtansine and Trastuzumab deruxtecan^3^, ADC development has primarily focused on three engineering axes: antigen selection, linker and payload optimization^4,5^. Despite substantial chemical and biological innovation, the clinical efficacy of ADCs remains constrained by the selective expression of tumor-associated antigens (TAAs), payload-related off-tumor toxicities, and resistance mechanisms^6–8^. The identification of highly specific TAAs continues to represent a major bottleneck, as high differential expression between tumor and normal tissues is considered advantageous for successful ADC development^1,9^. Likewise, linker optimization and payload design critically influence intracellular drug release and bystander effects, thereby improving the efficacy and safety profile of ADCs^2,10^.

A central determinant of ADC efficacy is the internalization of the antibody–drug conjugate following antigen binding. Importantly, internalization is not a simple extension of binding affinity to the target antigen. Although antibody discovery pipelines prioritize high-affinity clones against tumor-associated antigens (TAAs), efficient intracellular trafficking strongly depends on a complex interplay of multivalency, receptor clustering, membrane organization, and interfacial energetics that are largely dictated by the properties of the target receptor^11^. Clinical experience illustrates this distinction. Antibodies directed against HER2, such as trastuzumab, bind with high affinity yet induce relatively slow receptor internalization and substantial recycling^12^. HER2-directed ADCs including trastuzumab emtansine and trastuzumab deruxtecan achieve clinical efficacy despite these intrinsic trafficking constraints, relying on optimized linker chemistry, drug-to-antibody ratio (DAR), and payload permeability to compensate for moderate cellular uptake^13,14^. Conversely, ADCs such as sacituzumab govitecan leverage high receptor density and multivalent engagement to promote clustering-dependent endocytosis^15^.

A critical limitation in current clonal selection strategies is the reliance on static affinity metrics, which poorly predict whether a given antibody will be internalized after receptor engagement^16^. Efficient intracellular delivery requires that antibodies not only bind their target antigen but also induce productive conformational rearrangements that facilitate endocytosis^17^. Mapping the dynamic interfacial energy landscape of antibody– antigen complexes provides a route to identify clones predisposed to generate favorable contact networks and mechanically stimulate internalization. Antibody multivalency introduces an additional level of complexity: each heavy and light chain can engage antigen epitopes in multiple geometries, creating diverse binding modes that differentially modulate receptor organization and downstream trafficking. Such multivalent interactions can enhance avidity, promote cooperative receptor clustering, and generate energetically distinct interfaces that govern internalization kinetics^11,12,18^. Importantly, the impact of these interactions also depends on the receptor family involved. Distinct receptor classes—including receptor tyrosine kinases, immune receptors, and G-protein–coupled receptors—possess different internalization motifs and trafficking pathways that influence how antibody-induced clustering translates into endocytosis and intracellular routing^19^. Capturing these effects therefore requires a dynamic, ensemble-based framework that accounts for both the conformational plasticity of the antibody and the topology of the antigenic surface.

At the molecular level, antibodies and ADCs behave as dynamic conformational ensembles rather than static binding entities^20,21^. Sequence variation, chemical conjugation, and environmental context reshape conformational distributions and interfacial electrostatics, with direct consequences for tumor targeting and intracellular trafficking^9^. Hydrophobic payloads can further modulate membrane interactions and cellular uptake, while simultaneously increasing the risk of aggregation and off-target effects^22,23^. In that respect, computational biophysics approaches, including atomistic simulations and chemically accurate coarse-grained models^24–26^, provide a framework to resolve these dynamic and multiscale phenomena beyond sequence-centric or static structural descriptions.

Here we propose that antibody internalization can be rationalized by reconstructing the binding energy landscape governing antibody–receptor engagement. Using atomistic molecular dynamics (MD) simulations, we analyze a set of newly generated monoclonal antibodies targeting Junctional adhesion molecule A (JAM-A), a tight junction-associated receptor dysregulated in epithelial malignancies^27^. JAM-A is a transmembrane protein composed of a single transmembrane helix and two extracellular Ig-like domains that mediate homophilic and heterophilic interactions at cell–cell junctions^28,29^. These extracellular domains constitute the primary recognition sites for antibody binding, making JAM-A a suitable target for selective antibody-mediated delivery in tumor cells^30^. Rather than ranking mAb clones solely by binding affinity, we map intermolecular contact fingerprints, identify dominant energetic hubs, and resolve receptor–receptor interaction networks emerging upon antibody binding. We show that internalizing and non-internalizing clones exhibit distinct interfacial topologies: internalizing antibodies are more labile and display multivalent binding modes that may promote cooperative receptor clustering, thereby lowering the effective energetic barriers associated with early endocytic events.

Building on this framework, we demonstrate that MD-guided clonal selection provides a predictive strategy for engineering tumor-internalizing antibodies and optimizing next-generation ADCs. Our approach integrates sequence analysis, structure prediction using ColabFold^31–35^, and atomistic MD simulations of antibody–antigen complexes. Binding free energies are quantified using Umbrella Sampling^36^, enabling residue-level energetic decomposition and quantitative scoring of antibody clones. These computational predictions are directly validated through immunofluorescence-based internalization assays in tumor cells, establishing a correlation between binding modes, their associated energetics, and cellular uptake. Finally, we explore multivalent binding configurations and propose a mechanistic model linking dynamic binding behavior to antibody internalization.

## II. RESULTS AND DISCUSSIONS

### A. An MD-based computational framework for clonal selection

To establish a physics-based framework for mAb evaluation, we implemented a multistep computational pipeline integrating sequence analysis, structural prediction, and all-atom MD simulations. We first analyzed the sequences of rabbit mAbs targeting JAM-A classifying residues according to their physicochemical properties (Fig. 1A). These studies revealed distinct distributions of charged *vs*. hydrophobic amino acids within the complementarity-determining regions (CDRs), suggesting potential differences between the binding topology of the different clones. This sequence-level analysis provided us the initial basis for anticipating functional diversity beyond binding affinity measurements alone.

**FIG. 1.**
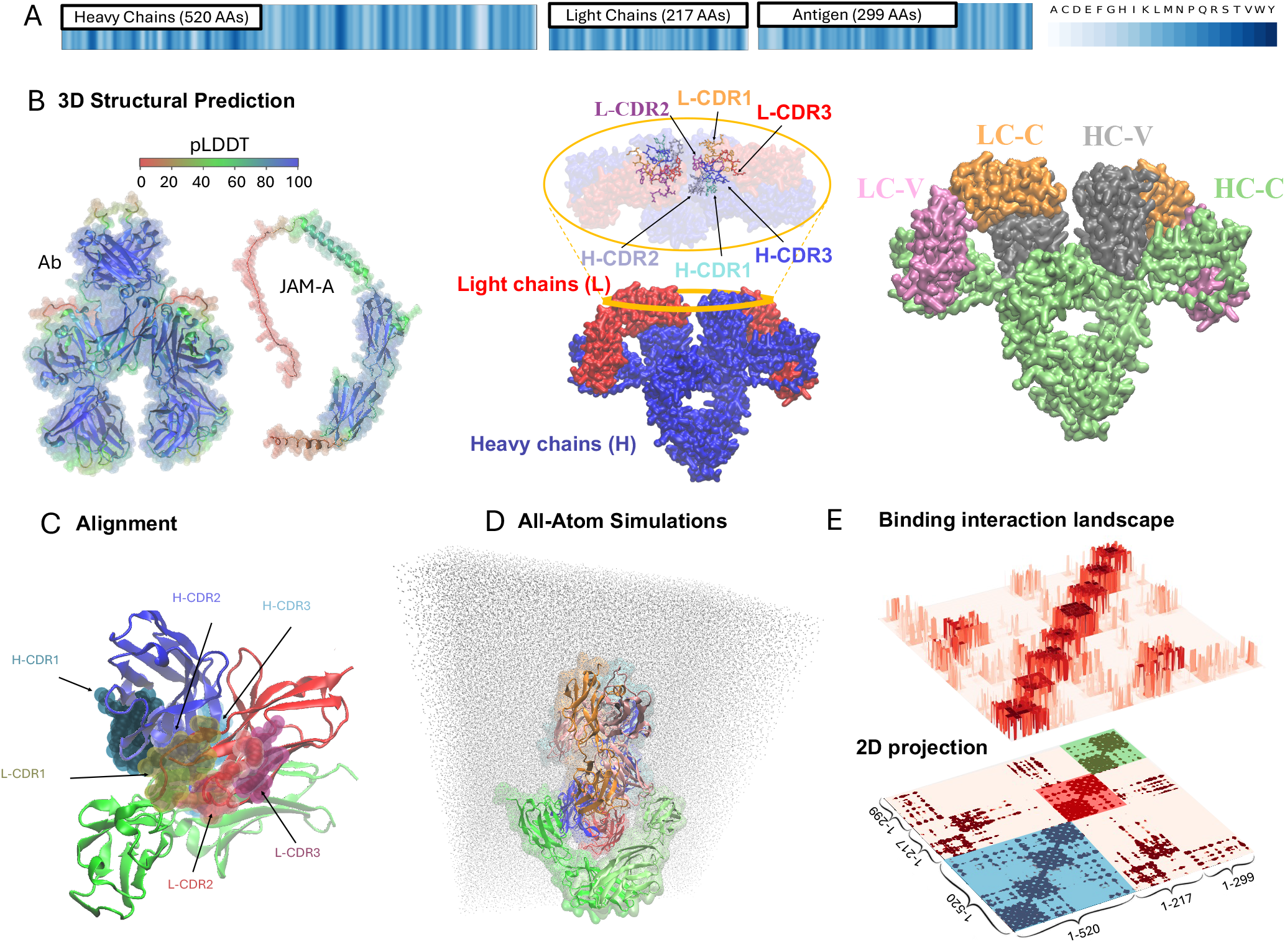
Structural and computational framework of the anti–JAM-A antibody system. A. Amino acid sequences of the monoclonal antibody heavy and light chains together with the target antigen JAM-A. Residues are color-coded according to physicochemical class, as indicated in the legend, highlighting the distribution of polar, charged, hydrophobic, and special-function amino acids across antibody and antigen sequences. B. Predicted molecular models obtained using ColabFold^31–35^. Left, structural models of the antibody and the JAM-A ectodomain. Structures are colored according to per-residue predicted Local Distance Difference Test (pLDDT) confidence scores, with the color gradient indicating model reliability at atomic resolution. Center, antibody representation highlighting heavy and light chains, together with a top-view inset showing the spatial localization of the complementarity-determining regions (CDRs). Right, antibody domains colored to distinguish variable and constant regions of the heavy and light chains. C. Identification of complementarity-determining regions (CDRs). The three heavy-chain CDRs are shown in blue and the three light-chain CDRs in red. The predicted antigen-binding interface is displayed in green, illustrating the spatial convergence of CDR loops toward the epitope region. D. Global view of the simulated system. The antibody is shown with its two heavy chains (blue and cyan) and two light chains (red and orange) in complex with two JAM-A molecules (green). The full assembly represents the starting configuration for molecular dynamics simulations. E. Binding interaction landscape of the antibody–antigen complex. The upper panel shows the interaction energy surface, whereas the lower panel displays its two-dimensional projection, which facilitates visualization of the corresponding contact maps. Blue, red, and green boxes highlight representative intramolecular interactions between antibody heavy chains, antibody light chains, and antigen–antigen contacts, respectively.

Subsequently, high-confidence structural models were generated. The predicted antibody structures displayed well-resolved ordered regions and flexible CDR loops, with per-residue predicted Local Distance Difference Test (pLDDT) scores indicating high structural reliability across the variable antibody domains. Structural models of the antibody and JAM-A ectodomain reveal the organization of heavy and light chains, the spatial localization of the CDRs, and the distribution of variable and constant domains that define the antigen recognition interface (Fig. 1B). In parallel, the JAM-A model was generated, exhibiting a stable *β*-strand-rich extracellular domain consistent with immunoglobulin-like folds. Importantly, the predicted antibody–antigen complexes revealed spatial convergence of the six CDR loops toward a defined epitope region, establishing the structural context for intermolecular binding analysis. Although static structural prediction provides a geometrically optimized initial binding configuration (Fig. 1C), it does not capture conformational plasticity or transient intermolecular interactions. Therefore, we performed MD simulations of the antibody–JAM-A complex with the all-atom Amber99sb force field^37^ in explicit solvent to resolve the time-dependent dynamics of the intermolecular binding (Fig. 1D). These simulations enabled the quantification of: (i) persistent antibody–antigen contacts, (ii) intra- and intermolecular interaction networks, and (iii) the dynamic inter-domain rearrangements controlling the interfacial binding energy landscape. In addition, a two-dimensional projection of the binding landscape was generated to facilitate visualization of the corresponding contact patterns (Fig. 1E).

### B. Binding free energy evaluation to predict mAb internalization

To quantitatively evaluate the thermodynamic landscape governing antibody–antigen recognition, we performed Umbrella Sampling simulations for five anti– JAM-A mAb clones (4F12, 5G11, 25C7, 25F9, and 22G6). For each clone, the Fab was simulated at increasing separation distances from the ectodomain of JAM-A, and the resulting trajectories were used to reconstruct the potential of mean force (PMF) profile (Fig. 2A). The reconstructed free-energy profiles reveal substantially different binding pathways among clones. Although all complexes exhibit a defined energetic minimum corresponding to the bound state, the depth, curvature, and width of the PMF profiles differ markedly. Clones displaying the deepest energy well do not necessarily follow identical association/dissociation pathways, indicating that binding strength alone does not capture the full complexity of the binding stabilization mechanisms. A difference in binding free energy of this magnitude has profound implications for complex stability and function. A PMF with ΔG ≈ −40 kJ mol^−1^ corresponds to a highly stable complex, in which the antibody–antigen interaction is strongly favored and dissociation events are rare on biologically relevant timescales. In contrast, a ΔG≈ −18 kJ mol^−1^ reflects a much weaker interaction, characterized by frequent association–dissociation events and a highly dynamic binding equilibrium.

**FIG. 2.**
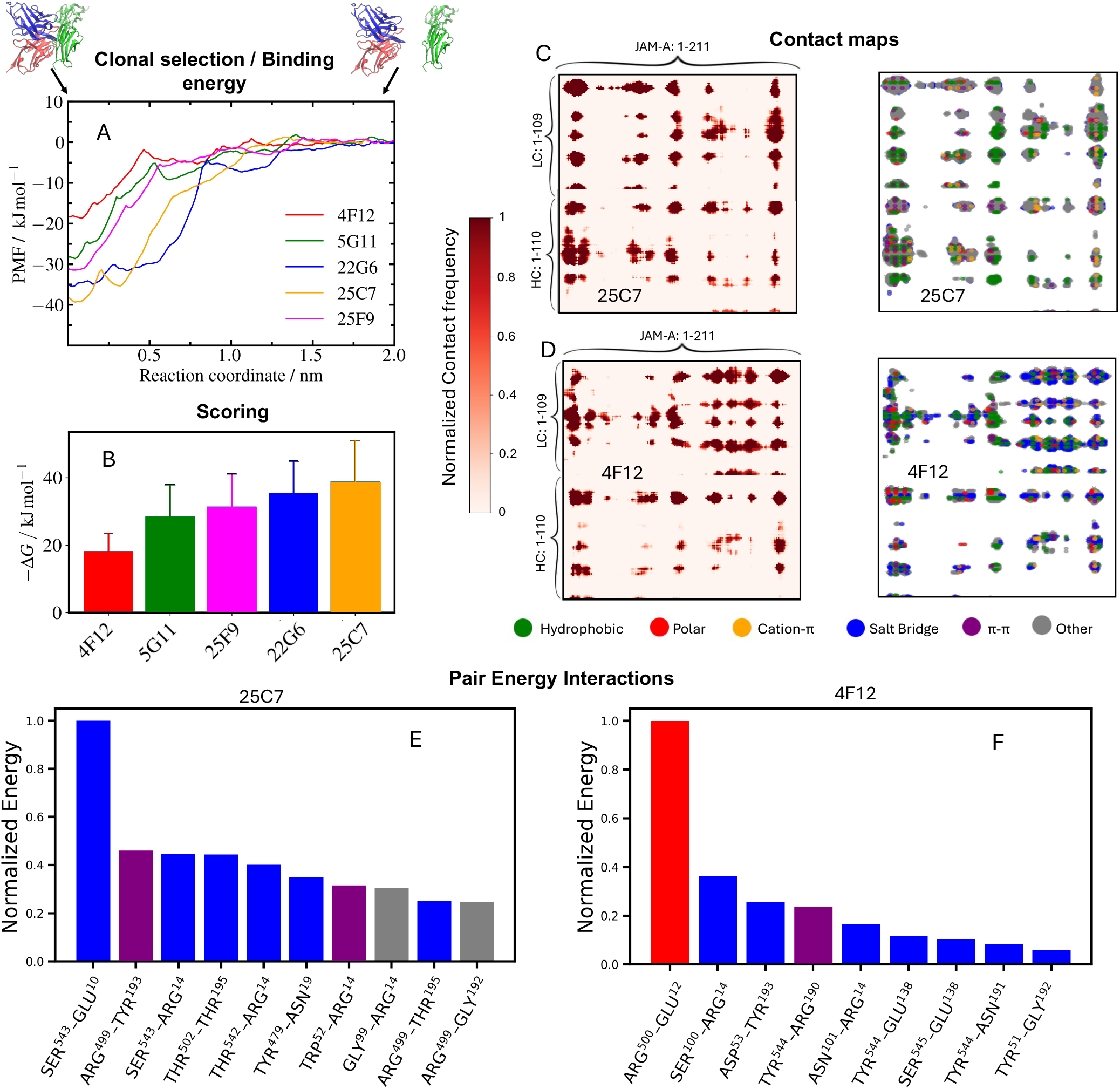
Free-energy landscapes and interfacial interaction patterns discriminate antibody clones. A) Potentials of mean force (PMFs) obtained from umbrella sampling simulations for the five antibody clones. The reaction coordinate corresponds to the center-of-mass distance between the Fab fragment and the JAM-A antigen. Representative simulation snapshots illustrating the associated (bound) and dissociated states of the complex are shown above the profiles. B. Binding free energies extracted from the PMFs for each clone and ranked from lowest to highest affinity. C–D. Intermolecular contact maps for the clones 25C7(C) and 4F12 (D). For each clone, the left panels show the contact frequency between antibody and antigen residues, while the right panels classify contacts according to their interaction type. Differences are particularly evident in the interaction patterns involving the variable region of the Fab light chain and distinct regions of the JAM-A receptor. E–F. Pairwise interaction energies for the ten most favorable intermolecular residue pairs at the antibody–antigen interface for clones 25C7 (E) and 4F12 (F), highlighting the emergence of dominant energetic hotspots in the high-affinity clone compared with the more distributed interaction network observed in 25C7.

From the PMF profiles, we derived binding free-energy estimates that enabled quantitative scoring of each clone (Fig. 2B). To further dissect the molecular determinants underlying these binding affinity differences, we analyzed the intermolecular contact frequencies through a distance cut-off analysis^38^ between the antibodies and JAM-A residues exhibiting the lowest- and highest-affinity, i.e., 25C7 and 4F12 clones, respectively. The resulting contact maps (Fig. 2C–D) reveal pronounced differences in the interaction patterns established between the Fab light chains and the JAM-A receptor. In particular, contacts involving the variable region of the light chain (LC:1–109) displayed markedly different distributions along the JAM-A sequence. Two major regions of the antigen can be distinguished (JAM-A:1–100 and JAM-A:101–211), where the overall contact density decreases for the high-affinity clone and reorganizes into a distinct interaction landscape for the lowest-affinity clone. Beyond these structural differences, analysis of interaction types revealed a striking shift in the physicochemical nature of the antibody–antigen interface. For the 25C7 clone, the contact maps show a dominant interaction type of hydrophobic-driven intermolecular contacts (Fig. 2C-right), complemented by a heterogeneous network of contacts enriched in cation-*π* interactions and polar residues. In contrast, the interaction pattern observed for the 4F12 clone (Fig. 2D-right) is characterized by a clear enrichment of polar contacts and electrostically-driven interactions, indicating a more specific and localized stabilization of the binding interface.

To quantify the energetic contribution of individual contacts, we calculated pairwise interaction energies for the ten most favorable residue pairs at the interface (Fig. 2E–F). For the 25C7 clone, the strongest interaction corresponds to a polar contact between serine^Ab^ and glutamic acid^JAM-A^, followed by cation-*π* contacts such as arginine^Ab^-tyrosine^JAM-A^ and arginine^Ab^-tryptophan^JAM-A^, as well as polar-like contacts mediated by serine, threonine, asparagine and glycine. For the 4F12 clone, the strongest interaction corresponds to an electrostatic contact between arginine^Ab^ and glutamic acid^JAM-A^. Additional energetic contacts are mediated by residues including tyrosine, serine, asparagine and glycine. The differences between the most energetic contact pairs and their location across the sequence for each clone highlight the presence of concentrated binding hotspots which modulate the global binding free energy as well as the binding association pathway between the CDRs and the epitope.

To experimentally evaluate antibody internalization, we performed immunofluorescence assays in ovarian cancer cells (OVCAR-3) using the different mAb clones (Fig. 3). Cells were treated with the antibodies for 0 and 12 hours, as described in the Materials and Methods section. As shown in Fig. 3A, the clones displaying clear internalization after 12 hours were 5G11 and 4F12, consistent with the computational predictions exhibiting lower binding free energy. Remarkably at 0 hours the strongest membrane-associated signal was observed for clones 25F9, 25C7, and 22G6, which also matches those with the highest calculated binding affinities (Fig. 2B). Internalization percentages were quantified as described in the Materials and Methods section. As shown in Fig. 3B, clones 4F12 and 5G11 exhibit significantly higher internalization levels than clones 25F9, 25C7, and 22G6. In this panel, the internalization percentages are compared against the binding free energies obtain in simulations, highlighting a well-defined energetic regime associated with internalization between binding interactions and cellular uptake Notably, clones with the lowest calculated free energies—reflecting the highest predicted affinities—did not correlate with productive cellular up-take. This observation suggests the existence of an optimal binding interaction energy range for internalization. Excessively strong binding may kinetically stabilize receptor engagement, potentially limiting the dynamic rearrangements required for receptor clustering and endocytosis, whereas moderate and multivalent binding energies may allow sufficient conformational plasticity to enable productive intracellular trafficking.

**FIG. 3.**
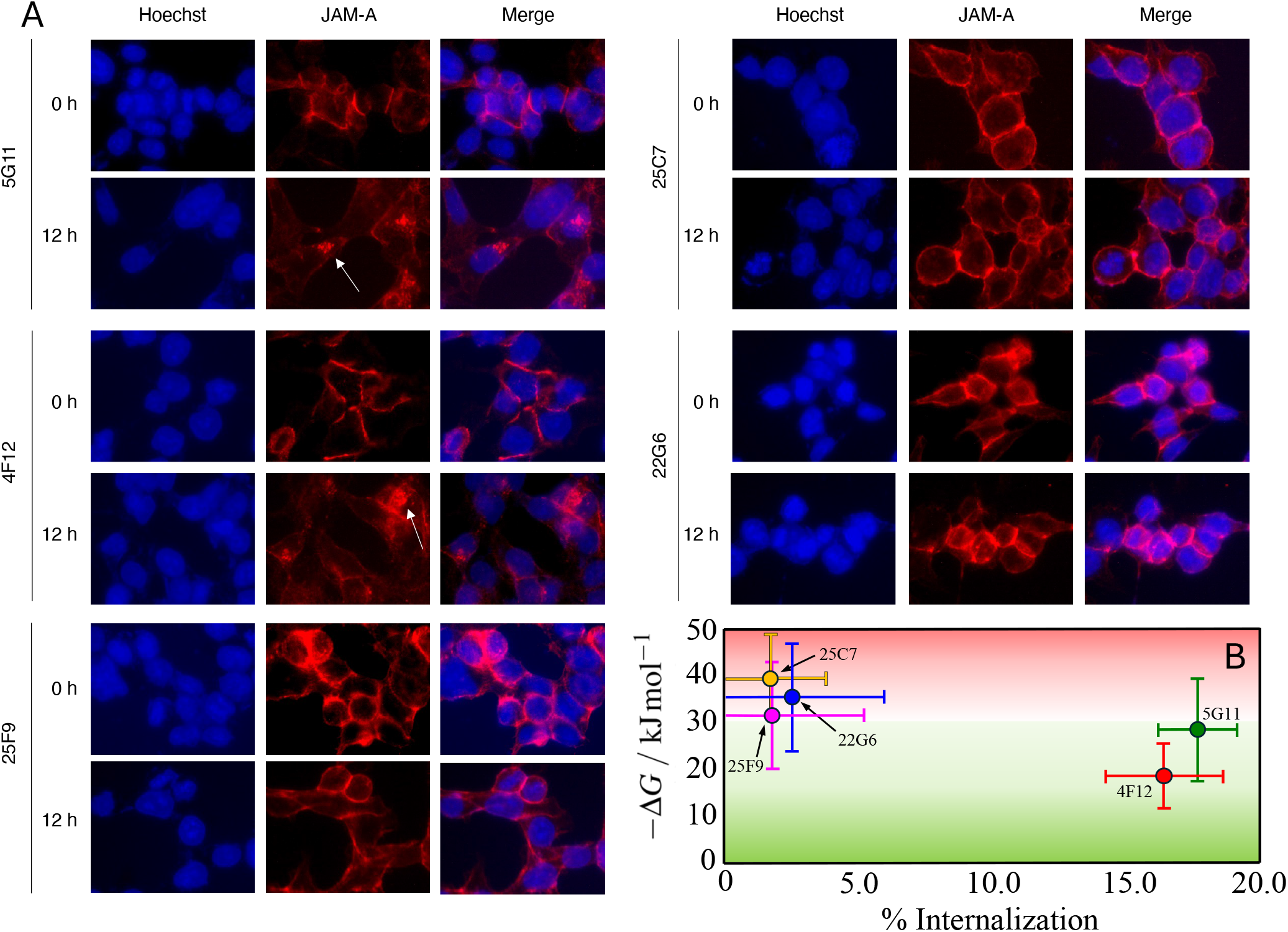
A. Evaluation of the internalization of the five monoclonal antibodies against the JAM-A protein. Treatments were carried out for 0 hours (30 minutes of incubation with each antibody at 37ºC followed by fixation) and 12 hours (treatment for 30 minutes, following by washing and incubation for 12 hours at 37ºC). White arrows indicate intracellular accumulations of the antibody signal. Inmunofluorescence was then performed as indicated in methods. Images were taken using a Nikon Eclipse E200 inverted fluorescence microscope and processed using ImageJ software. Computed binding free energy plotted as a function of the experimentally measured percentage of antibody internalization. Each point represents an antibody variant. Shaded regions red and green indicate energetic internalization regimes incompatible or compatible respectively.

### C. Mechanistic analysis of multivalent binding modes and antibody internalization

Next, we studied a humanized antibody derived from the rabbit clone 4F12 and investigated the mechanistic basis of its internalization. Although JAM-A is over-expressed in ovarian tumor cells, it normally functions as a tight-junction protein, with its spatial organization tightly regulated in epithelial tissues^28^. In tumor cells, disruption of this architecture leads to a more heterogeneous distribution of JAM-A across the plasma membrane, which can limit the local receptor density accessible to antibodies^39^. Given this heterogeneous distribution, the mode of antibody engagement—whether monovalent or bivalent—may critically influence internalization. We explored possible modes of engagement by considering the stoichiometry of binding: i) a monovalent interaction involving one Fab and one JAM-A receptor (1 Fab–1 receptor), ii) a partially bivalent complex in which both Fab domains are present but only one receptor is engaged (2 Fab–1 receptor), and iii) a fully bivalent complex engaging both Fab domains with two receptors (2 Fab–2 receptor). To dissect the internalization mechanism of the humanized 4F12 antibody, we analyzed the binding energetics associated with each stoichiometric scenario. Binding free energies were estimated using umbrella sampling simulations, and PMFs (Fig. 4B) were reconstructed along a normalized reaction coordinate, with 0 representing maximum contact and 1 representing distances at which their interaction vanishes into zero. Such normalization allowed direct comparison between the different binding modes despite variations in absolute center-of-mass distances.

**FIG. 4.**
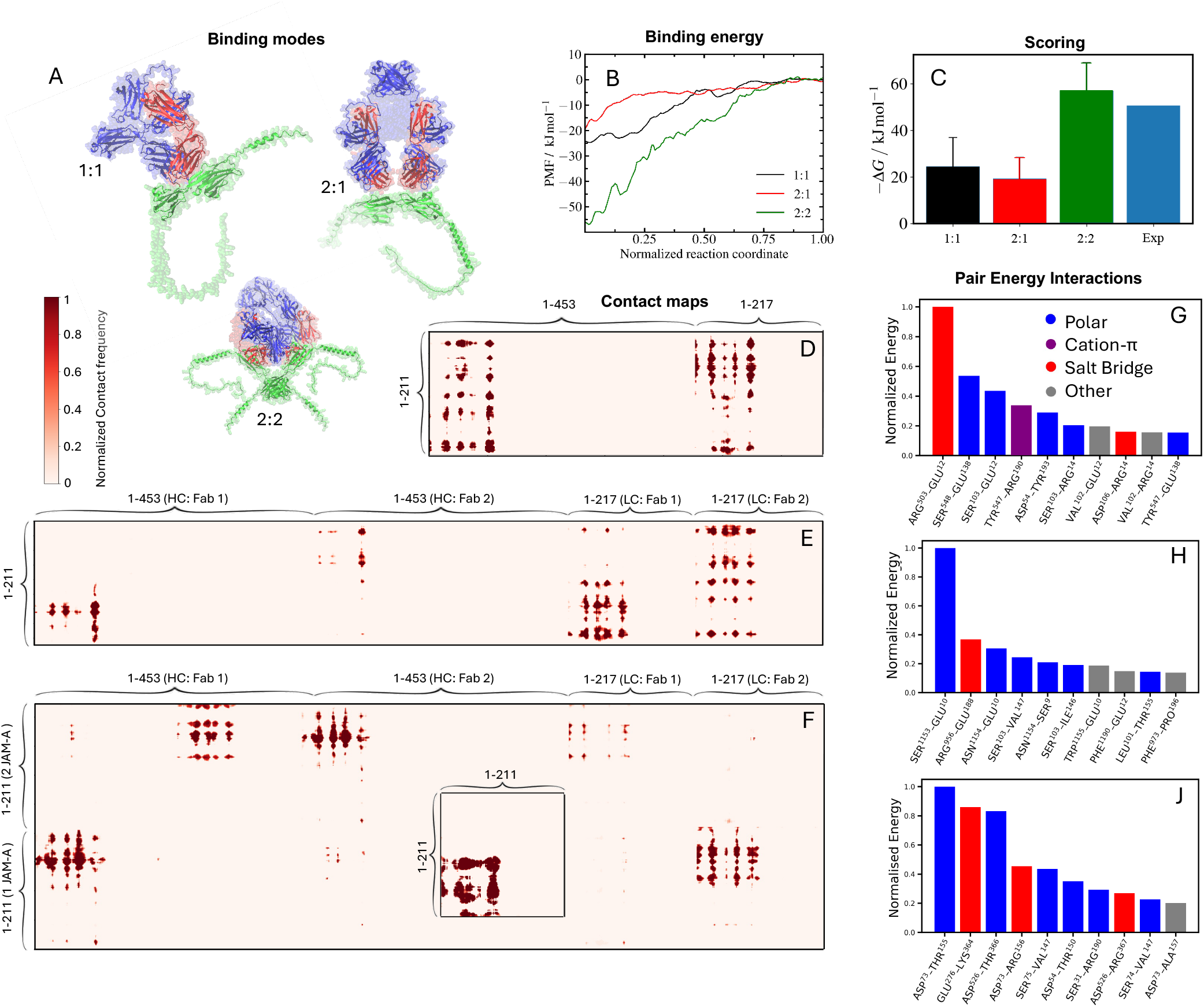
Mechanistic characterization of humanized 4F12 binding and internalization. A. Structural visualization of the three predicted binding modes I: 1:1, II: 2:1 and III: 2:2. The JAM-A antigen(s) are shown in green, while the antibody heavy and light chains are depicted in blue and red, respectively. B. PMFs computed for the three binding modes. C. Quantitative scoring of each mode based on ΔG extracted from PMFs, compared with experimental SPR measurements. D–F. Residue-residue contact maps for modes I, II, and III, respectively; F inset highlights contacts between the two JAM-A antigens in the III mode. G–J. Interaction energies of the ten most stabilizing residue pairs for modes I, II, and III, respectively.

To dissect the internalization mechanism of the humanized 4F12 antibody, we analyzed the binding energetics (Fig. 4C) and residue-residue interactions of the three principal binding modes (Fig. 4D–J). The initial recognition mode (mode I: 1 Fab–1 receptor) exhibits a moderate binding free energy of approximately 23 kJ mol^−1^. The subsequent interaction mode (mode II: 2 Fab–1 receptor) shows a similar energy of 20 kJ mol^−1^, suggesting that these two binding modes may coexist in a highly dynamic equilibrium across lateral diffusion over the cell surface. This multivalent binding mode may allow the antibody–receptor complex to diffuse and efficiently encounter a second JAM-A receptor, ultimately enabling bivalent engagement.

Analysis of the contact maps highlights the multivalent behavior of the antibody. In mode II, each Fab engages a distinct Ig-like domain of the single JAM-A antigen, preventing symmetric contacts. In contrast, the fully bivalent 2:2 mode shows increased avidity and partially symmetric cooperative interactions. The first JAM-A antigen primarily engages the variable region of the heavy chain of Fab 1, with weaker interactions to Fab 2, while also engaging the variable region of the light chain of Fab 2. The second JAM-A antigen interacts strongly with the light chain of Fab 2 and with the constant region of the heavy chain of Fab 1. Contacts with the light chain of Fab 1 are weaker, but overall the pattern demonstrates symmetry across the two Fab domains (Fig. 4D-F). Additionally, the two JAM-A antigens interact via their first Ig-like domains, further stabilizing the multivalent complex (Fig. 4F; inset). These maps illustrate how multivalency distributes binding across multiple interfaces, enhancing avidity and stabilizing the fully engaged complex.

Once the second receptor is engaged, the system reaches the fully bivalent “internalization binding mode” (mode III: 2 Fab–2 receptor), forming a trimeric complex composed of two JAM-A antigens and one antibody. This state exhibits a substantially higher binding free energy of 58 kJ mol^−1^, stabilizing the complex at the membrane and enabling the initiation of membrane curvature required for early endocytosis. The high affinity is mechanistically necessary, as bending the membrane incurs an energetic penalty proportional to the bending modulus and interfacial rigidity, reflecting the excess area generated during curvature. To validate these computational predictions, we also performed surface plasmon resonance (SPR) experiments of the humanized 4F12 clone to measure the antibody’s affinity for JAM-A. Remarkably, the experimental SPR value (51 kJ mol^−1^; Fig. 4C) near-quantitatively matches the simulation prediction, reinforcing the hypothesis that the 2:2 complex represents the energetically preferred and functionally relevant binding mode. By achieving sufficient interaction energy, the antibody–receptor complex can overcome the mechanical cost of membrane curvature, effectively anchoring to the membrane and promoting early endocytosis.

An important mechanistic implication of our results is that both the orientation and the affinity of the antibody–antigen interaction critically determine productive internalization. In the context of tight junctions, where JAM-A molecules are presented at cell–cell interfaces, the antibody may initially bind receptors from two opposing cells. If this bivalent configuration is associated with excessively high affinity, the antibody–receptor complex may become kinetically trapped, preventing the reorganization required to engage receptors on the same membrane and transition toward the internalization-competent state. Thus, the lability of the initial binding mode is essential to avoid such kinetic arrest, enabling the antibody to dissociate and rebind in alternative configurations that favor productive multivalent engagement and subsequent endocytosis. A schematic representation of this multi-step internalization mechanism is shown in Fig. 5.

**FIG. 5.**
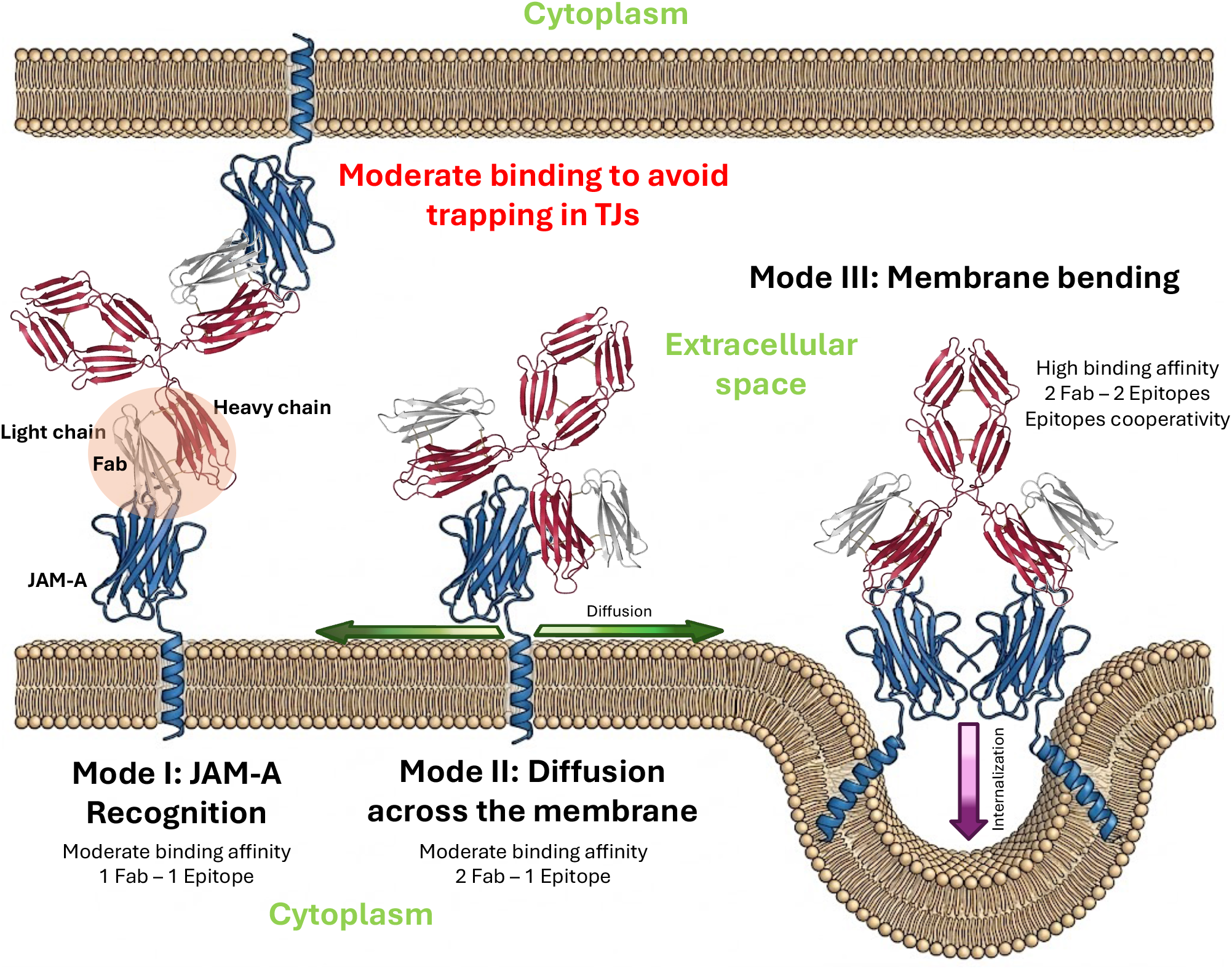
Schematic representation of the proposed internalization mechanism, illustrating the three binding modes and the antibody (red) sequential engagement with JAM-A receptors (blue) leading to membrane anchoring and pre-endosome formation.

Another observation emerging from our analysis concerns the nature of the 2:1 binding configuration. Although the mode II is consistently predicted by AlphaFold, we do not consider it to be a biologically relevant stable state. The interaction in this mode is intrinsically asymmetric between the two Fab domains, lacking the cooperative engagement that characterizes productive multivalent binding. Instead, we propose that this configuration represents a transient intermediate along the binding pathway. In this framework, the initial interaction is established through one Fab in mode I, which remains in a dynamic equilibrium between association and dissociation. From this state, the antibody can undergo reorientation events that allow the second Fab to engage the receptor. This transition may occur either through partial dissociation followed by rebinding, or through a direct rearrangement of the antibody while maintaining contact with the receptor. In this sense, mode II can be interpreted as a transition state connecting two equivalent configurations of mode I, Fab 1–receptor and Fab 2–receptor binding—facilitating configurational exploration prior to productive multivalent engagement. Notably, the simultaneous Fab–antigen/Fab–antigen interaction is equivalent to that described for mode III.

## III. CONCLUSIONS

In this study, we combined protein structural prediction, MD simulations, and *in vitro* cell-based immunofluorescence experiments to dissect the molecular determinants of antibody internalization in tumor cells. Using antibodies targeting JAM-A, a tight-junction-associated receptor dysregulated in epithelial malignancies, we demonstrated that moderate binding energies (i.e., between 20-30 kJ/mol) between one Fab domain and one receptor correlate with productive cellular uptake. Our all-atom PMF calculations, contact map analyses and energetic decomposition, combined with immunofluorescence assays suggest that moderate binding affinities enable dynamic exploration of the cell surface, facilitating further engagement with two receptors in a multivalent manner. Such stable engagement ultimately drives intracellular uptake.

Our fully humanized 4F12 antibody exhibits a hierarchy of binding modes, from initial recognition (1 Fab–1 receptor) to a bivalent, high-affinity internalization binding mode. Contact maps show that multivalency distributes interactions across multiple sequence interfaces, including antibody–antigen and antigen–antigen contacts, enhancing avidity and stabilizing the fully engaged complex. The high binding energy—evaluated through PMF calculations and SPR in vitro experiments—of the internalization mode (e.g., of 50-60 kJ mol^−1^) is mechanistically required to overcome the energetic cost of membrane bending during pre-endosome formation, linking molecular recognition directly to cellular mechanics. This framework highlights a mechanistic principle for antibody design that is well-characterized in other contexts but largely unexplored in ADC development^11^: achieving an optimal balance between affinity, multivalency, and dynamic receptor engagement is critical for efficient internalization. Excessive affinity in the initial recognition step may impede receptor search, dynamic orientational rearrangements to find the optimal pose, and limit productive uptake. In contrast, moderate binding affinities maintain dynamic attachment, thereby facilitating the formation of multivalent complexes formed by two receptors and one antibody, capable of anchoring and deforming the membrane.

Our integrated computational-experimental approach provides a quantitative platform for clonal selection and antibody engineering, enabling the identification of candidates that combine favorable binding energetics with significant internalization kinetics. Beyond JAM-A, this strategy is broadly applicable to the rational design of antibodies and ADCs against cell-surface receptors where internalization and receptor clustering are essential for therapeutic efficacy. In conclusion, our results define a principled pathway from sequence to mechanism, demonstrating that molecular dynamics-informed clonal selection can guide the design of antibodies with optimal cellular uptake. Thus, this study establishes a generalizable methodology for rational engineering of next-generation antibody therapeutics.

## IV. METHODS

### Simulation details

The JAM-A antigen structure was predicted using ColabFold^31–35^ and cut to retain only the extracellular Ig-like domains, removing flexible residues outside of the structured binding regions. The antibody was truncated to its Fab fragment, with Fc residues removed, ensuring that the simulations focused exclusively on the recognition interface. The Systems were solvated with TIP3P water and neutralized with counterions, followed by energy minimization. All simulations were performed in GROMACS^40^ with the AMBER99SB-ILDN force field^37^. Electrostatics were treated with PME^41^ and a 1.4 nm cutoff and van der Waals interactions were truncated at the same distance. Temperature and pressure were maintained at 298 K and 1 bar using a velocity-rescale^42^ thermostat and a C-rescale^43^ barostat. Steered molecular dynamics (SMD) was applied to generate initial unbinding pathways. A harmonic pulling potential (k = 1000 kJ mol^−^1 nm^−^2) was applied along the center-of-mass reaction coordinate at 0.01 nm ps^−^1. Umbrella sampling^44–46^ was performed along the normalized reaction coordinate (0 = close contact to 1 maximal separation), the resulting histograms of the restricted positions were analyzed using the Weighted Histogram Analysis Method^47^ (WHAM) to reconstruct the potential of mean force (PMF). Mode I (1 Fab–1 receptor) used 13 windows, mode II (2 Fab–1 receptor) 27 windows, and mode III (2 Fab–2 receptor) 34 windows, with 5 ns of production per window. Statistical uncertainties were estimated using a bootstrap analysis, in which time correlations were explicitly accounted for by evaluating the integrated autocorrelation time of the reaction coordinate in each window Residue-residue contacts were computed between C*α* atoms with a cutoff of 2.5 nm and analyzed for frequency and interaction type (polar, nonpolar, electrostatic). Pairwise interaction energies were calculated for the ten most stabilizing residue pairs per mode, identifying energetic hotspots and allowing correlation between interfacial architecture, computed binding affinities, and experimental observations.

### Experimental details

An ovarian cancer cell line was seeded on glass coverslips and, after 24 hours, treated with each monoclonal antibody (10 nM) for the indicated times, consisting of 30 minutes of antibody incubation at 37°C, followed by fixation for time 0, or washing and incubation for 12 hours. The coverslips were then washed and fixed in 2% paraformaldehyde for 30 minutes at room temperature. The coverslips were then washed and incubated with a quenching solution (NH_4_Cl) in PBS +/+. For permeabilization, they were rinsed with PBS +/+ containing 0.1% Triton X-100 and 0.2% BSA for 10 minutes three times and washed with PBS +/+ containing 0.2% BSA. All chemicals were purchased from Sigma Aldrich (St. Louis, MO). The coverslips were then incubated with anti-rabbit Cy3 (1:750, Thermofisher Scientific, USA) for 30 minutes. Hoechst 33342 (1:10,000, MedChemExpress, Monmouth Junction, NJ, USA) was added and washed with PBS +/+. The mounting medium was FluorSave^™^ reagent (Merck, Darmstadt, Germany). Fluorescence images of the cells were obtained using a Nikon Eclipse E200 inverted fluorescence microscope and analyzed using ImageJ. The internalization percentage was calculated by counting the cells with internalised JAM-A from a total of 150–200 nuclei per condition.

Binding affinity measurements were performed by surface plasmon resonance (SPR) by GenScript using a Biacore 8K system (Cytiva). All experiments were conducted at 25°C using HBS-EP+ buffer (pH 7.4) as running buffer, prepared from a 10× stock solution. A Series S Sensor Chip CM5 functionalized with anti-histidine antibodies was used for ligand capture. Regeneration of the sensor surface between cycles was achieved using 10 mM glycine-HCl (pH 1.5).

The His-tagged ligand (JAM1 HUMAN) was captured onto the sensor chip surface at a concentration of 0.5 *µ*g/mL, reaching approximately 7 response units (RU) after a 30 s injection. Humanized 4F12 antibody was diluted in running buffer and injected over the immobilized ligand at a concentration of 100 nM. The association phase was monitored for 120 s, followed by a dissociation phase of 180 s, at a constant flow rate of 30 *µ*L/min.

Reference subtraction was performed using a double-referencing method, including both a reference flow cell and blank buffer injections. Sensorgrams were processed and analyzed using Biacore 8K Evaluation Software (version 5.0). Kinetic parameters, including the association rate constant (ka), dissociation rate constant (kd), and equilibrium dissociation constant (KD), were determined by fitting the data to an appropriate binding model.

## V. ACKNOWLEDGEMENTS

P.L acknowledges funding from the European Union’s Horizon 2020 research and innovation program (grant agreement 101160499 to J. R. E). J. R. E. acknowledges funding from Emmanuel College, the University of Cambridge, the Ramón y Cajal fellowship (RYC2021-030937-I), the Spanish scientific plan and committee for research reference PID2022-136919NA-C33, and the European Research Council (ERC) under the European Union’s Horizon Europe research and innovation program (grant agreement no. 101160499). C.N.J and A.O., acknowledge funding from CRIS Cancer Foundation (AOF.C01CRIS and AOF.M01CRIS).

## VI. AUTHORS CONTRIBUTIONS STATEMENT

P.L., C.N.-J., A.O., and J.R. contributed to the conceptualization of the study. P.L. performed the simulations and C.N.-J. carried out the experiments. A.O. and J.R. were responsible for funding acquisition. A.P., J.R., and A.O. supervised the project. P.L. and C.N.-J. wrote the original draft of the manuscript. All authors contributed to the writing and revision of the final paper.

