## Supporting information for "Computational mapping of antibody-receptor energy landscapes to predict internalization"

[4)] *Centro de Investigación del Cáncer. CSIC and CIBERONC, 37007-Salamanca, Spain*

[5)] *Cátedra INTHEOS-START-CEU de Oncología de precisión, Facultad de Medicina, Universidad San Pablo-CEU, Montepríncipe, 28660 Boadilla del Monte, Madrid, Spain*

[6)] *C-Therapeutics, TMC3 Elix Park 1885 Old Spanish Trl, Houston, TX 77039 United States*

[7)] *Yusuf Hamied Department of Chemistry, University of Cambridge, Lensfield Road, Cambridge CB2 1EW, UK*

(Dated: 25 March 2026)

[a)]Electronic mail:

[b)]Electronic mail:;

†These authors contributed equally

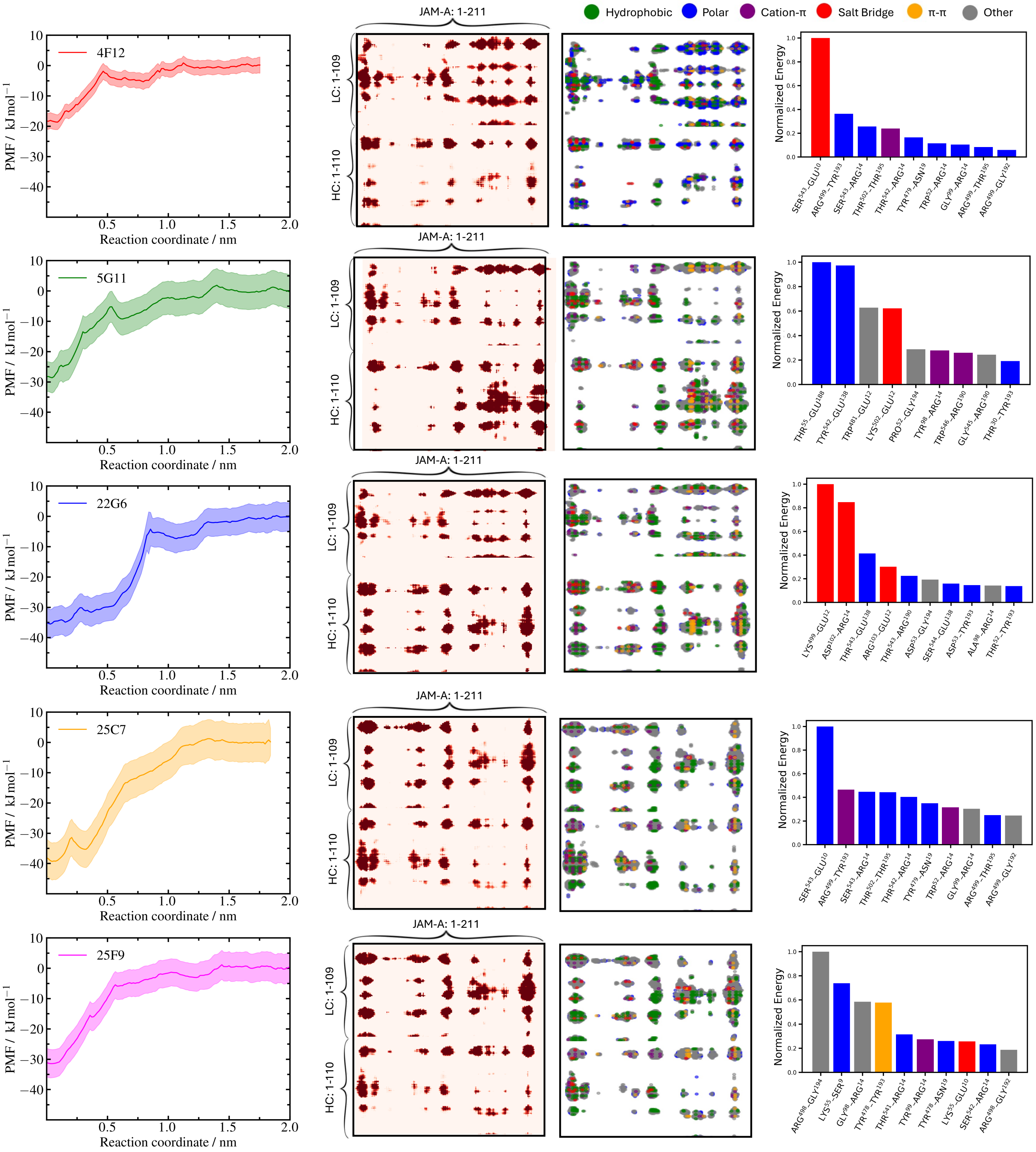


FIG. 1. Free-energy landscapes and interfacial interaction patterns discriminate antibody clones. Each row corresponds to one antibody clone. The left column shows the potentials of mean force (PMFs) obtained from umbrella sampling simulations. The reaction coordinate corresponds to the center-of-mass distance between the Fab fragment and the JAM-A antigen. Error bars were estimated by bootstrapping. The center-left column displays intermolecular contact maps normalized by contact frequency. The center-right column shows the same contacts classified according to interaction type, distinguishing the main physicochemical contributions at the antibody–antigen interface. The right column reports the ten most energetically favorable intermolecular residue pairs for each clone.

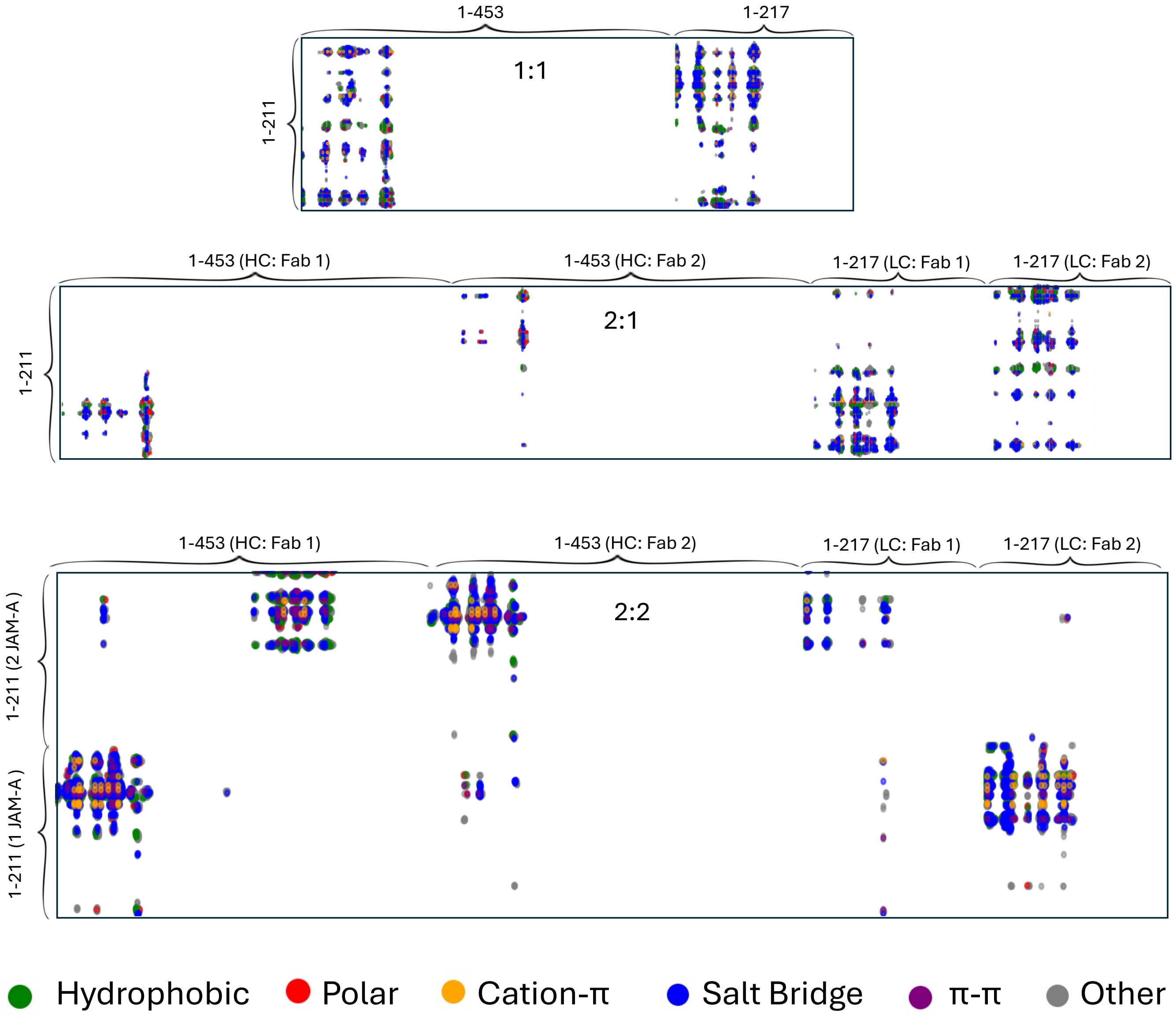


FIG. 2. Interaction-type contact maps for the different binding modes. The top panels show the intermolecular contact maps classified by interaction type for binding mode I (1 Fab–1 antigen). The middle panels correspond to the intermediate binding mode II (2 Fab–1 antigen), while the bottom panels represent the fully bivalent configuration (2 Fab–2 antigen). These maps highlight the distinct interaction patterns established at the antibody–antigen interface for each binding stoichiometry.
